# *Ocimum gratissimum* essential oil nanoemulsions as a safe topical nanoplatform for antibacterial and wound-healing activities

**DOI:** 10.64898/2026.07.01.735794

**Authors:** A Fomesseng Negoue, F Eya’ane Meva, J-B Hzounda Fokou, S Voundi Olugu, V Boudjeka, J C Ngo Nyobe, P Belle Ebanda Kedi, A M Houatchaing Kouemegne, G Etame Loe

**Affiliations:** Department of Pharmaceutical Sciences, Faculty of Medicine and Pharmaceutical Sciences, University of Douala, PO Box 2701, Douala, Cameroon; Biotechnology Laboratory, University Institute of Technology, University of Douala PO Box 8698, Douala, Cameroon

**Keywords:** *Ocimum gratissimum*, Essential oil, Nanoemulsion, Wound healing, Antibacterial activity

## Abstract

**Background:** Natural essential oils exhibit antimicrobial and wound-healing properties, but their therapeutic application is limited by poor water solubility, volatility, and instability. This study developed and characterized a nanoemulsion of *Ocimum gratissimum* essential oil (OGNe) and evaluated its physicochemical properties, dermal safety, antibacterial activity, and wound-healing potential.

**Methods:** Essential oil was obtained by hydrodistillation and formulated into nanoemulsions by high-speed stirring emulsification. Physicochemical properties, including pH, droplet size, polydispersity index, and storage stability, were determined. Acute dermal toxicity was assessed in Wistar rats following OECD Test Guideline 402. Antibacterial activity was evaluated using broth microdilution, minimum inhibitory concentration (MIC), minimum bactericidal concentration (MBC), and time-kill assays. Wound-healing efficacy was investigated using an excision wound model over 21 days using distilled water and trolamine serving as controls.

**Results:** OGNe exhibited a stable milky appearance, near-neutral pH, and droplet sizes ranging from 26 to 224 nm. No signs of dermal toxicity or behavioral abnormalities were observed after topical administration. The nanoemulsion showed selective antibacterial activity, with the highest susceptibility against *Acinetobacter baumannii* (MIC = 1.125 μL/mL), whereas *Escherichia coli* remained resistant. Time-kill assays demonstrated concentration-dependent bacteriostatic activity. In vivo, OGNe significantly accelerated wound contraction from day 3 onward (p < 0.0001), achieving healing rates comparable to or exceeding those of trolamine during the inflammatory and proliferative phases.

**Conclusion:** *Ocimum gratissimum* nanoemulsions represent stable, biocompatible topical formulations that combine selective antibacterial activity with enhanced wound healing, supporting their potential as phytopharmaceutical nanoformulations for the management of acute skin wounds.

## Introduction

A wound is defined as a disruption of the normal integrity of body tissues caused by physical injury or other damaging agents. Depending on the depth of tissue loss, wounds are classified as erosions, fissures, or ulcers, the latter extending into the dermis and frequently becoming chronic and difficult to heal. [1] The healing process depends on numerous factors, ranging from the patient’s general health status to the nature, severity, and treatment of the lesion. The pathophysiology of wound healing, its associated complications, and scar formation is multifactorial and highly complex. [1] Chronic wounds place a substantial burden on healthcare systems, particularly in resource-limited countries, because of their high prevalence and treatment costs. [2] Approximately 80% of the population in resource-limited countries relies on medicinal plants for primary healthcare because of their accessibility and affordability. *Ocimum gratissimum* L, popularly known as scent leaf, has long been used in traditional medicine for the management of diabetes, cancer, inflammation, anemia, diarrhea, pain, and fungal and bacterial infections. [3–5] Phytochemical investigations have shown that *O. gratissimum* essential oil contains camphene, β-caryophyllene, α- and β-pinene, α-humulene, sabinene, β-myrcene, limonene, 1,8-cineole, trans-β-ocimene, linalool, α- and δ-terpineol, eugenol, α-copaene, β-elemene, p-cymene, thymol, and carvacrol. [5–8] Silva et al. demonstrated, using 6–50% DMSO solutions of *O. gratissimum* essential oil, that neither the complete essential oil nor eugenol produced inhibition zones against the bacterial strains tested. [6] Nevertheless, combination assays with amikacin and erythromycin revealed synergistic interactions between *O. gratissimum* essential oil or eugenol and these antibiotics against *Escherichia coli* and *Staphylococcus aureus*. Conversely, gentamicin combined with *O. gratissimum* exhibited antagonistic activity against *S. aureus* but synergistic effects against *E. coli*. [6] The limited recovery of essential oils following extraction, together with their volatility and chemical instability under environmental conditions, necessitates the development of systems capable of improving the stability, protection, and bioavailability of their bioactive constituents. To overcome these limitations, several nanotechnology-based strategies have recently been explored. Our research group previously developed antimicrobial wound dressings by impregnating cotton fibers with silver nanoparticles and characterizing them using hyperspectral microscopy. [9] The wound-healing activities of *Diospyros hoyleana* extracts and green-synthesized silver nanoparticles derived from *Vernonia conferta* were successfully demonstrated in experimental rats. [8,10] Likewise, chitosan capsules encapsulating silver nanoparticles synthesized from *Elaeis guineensis* leaf extracts accelerated wound repair, whereas similar polymeric systems containing *Strychnos phaeotricha* bark extracts exhibited antimicrobial activity against *Salmonella* spp., *Escherichia coli*, and *Candida* spp. [11,12] Several investigations have demonstrated that the bioactive components contained in essential oils attach to microbial cell surface and penetrate the phospholipid bilayer of the cell membrane, followed by membrane damage, which causes negative impacts on cell metabolic activities and cell death. [13] Numerous approaches for the manufacturing essential oil-loaded nanoparticles including co-precipitation, high-pressure homogenization, high-speed stirring, ultra-sonication, ionic gelation, mini-emulsion polymerization, nanoprecipitation, spray drying, and the Stöber process have been developed. [13] Among these approaches, high-speed stirring provides a simple, scalable, and cost-effective strategy for producing stable nanoemulsions with reduced droplet size and increased interfacial surface area, properties that may enhance the biological performance of encapsulated essential oils. Therefore, the present study aimed to formulate a nanoemulsion of *O. gratissimum* essential oil using high-speed stirring and to investigate its antibacterial and wound-healing activities in experimental models.

## Experimental

### Collection, authentication, and preparation of extract

Fresh leaves of *O. gratissimun* (Figure 1) were harvested in Bafou (Dschang, Cameroon) between 5:30 and 7:00 a.m. Plant material was identified at the National Herbarium of Cameroon by comparison with voucher specimen Westphal No. 18741, and a voucher specimen No. 42738/SRF/Cam. Essential oil was extracted from fresh leaves according to the French Association for Standardization (AFNOR) guidelines using a Clevenger-type hydrodistillation apparatus. During distillation, the essential oil-saturated vapor condensed in the cooling system, producing two immiscible phases: an aqueous phase (hydrosol) and an upper organic phase containing the essential oil. The essential oil was recovered by decantation, stored in amber glass bottles, and dried over anhydrous sodium sulfate (Na₂SO₄) before further analyses.

**Figure 1.**
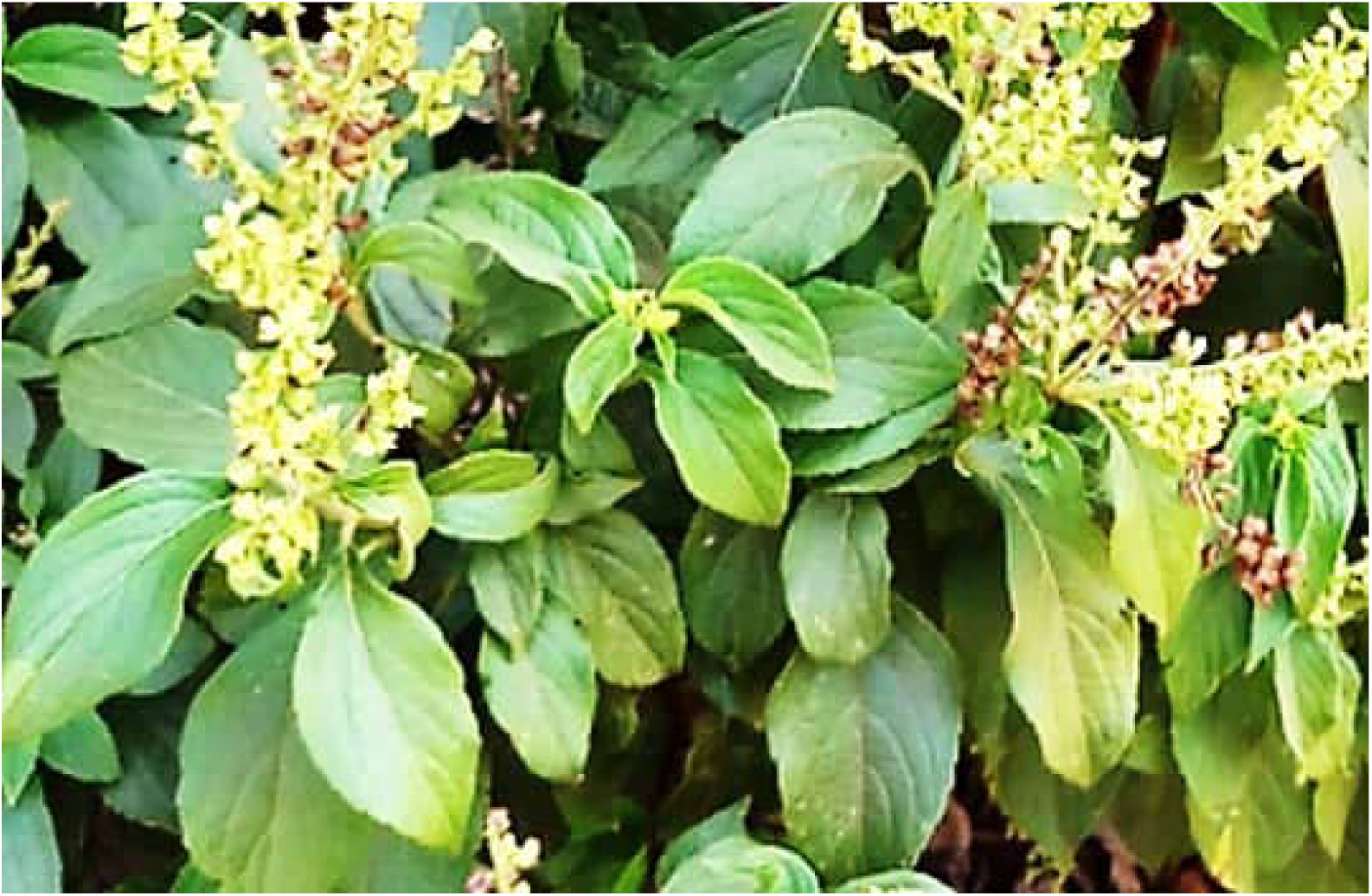
*Ocimum gratissimun* plant

The essential oil yield was estimated by the ratio of the masses of EO and fresh plant material and expressed as a percentage (%) (1).

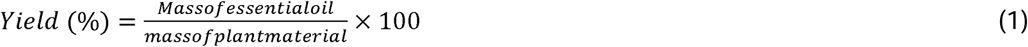

### Animal materials

Albino Wistar rats (*Rattus norvegicus*) were obtained from the animal facility of the Faculty of Medicine and Pharmaceutical Sciences, University of Douala. The study included 34 female rats aged between 8 and 10 weeks. Animals had free access to food and water throughout the experiment.

### Preparation of the nanoemulsion of *Ocimum gratissimum*

The essential oil nano-emulsion was prepared according to the method described by Purkait et al. (2020). [15] with minor modifications. *O. gratissimum* essential oil and Tween 80 were used as the organic phase (oil and surfactant) with a 1:1 ratio (v/v), while demineralized water constituted the aqueous phase. The essential oil and Tween 80 were mixed using a magnetic stirrer (magnetic stirrer hot plate 85-2, USA) at a speed of 500 rpm for 30 min. Subsequently, deionized water was gradually added under continuous stirring for 4 hours at room temperature followed by 1 h of sonication. The formulation composition is summarized in Table 1.

**Table 1.**
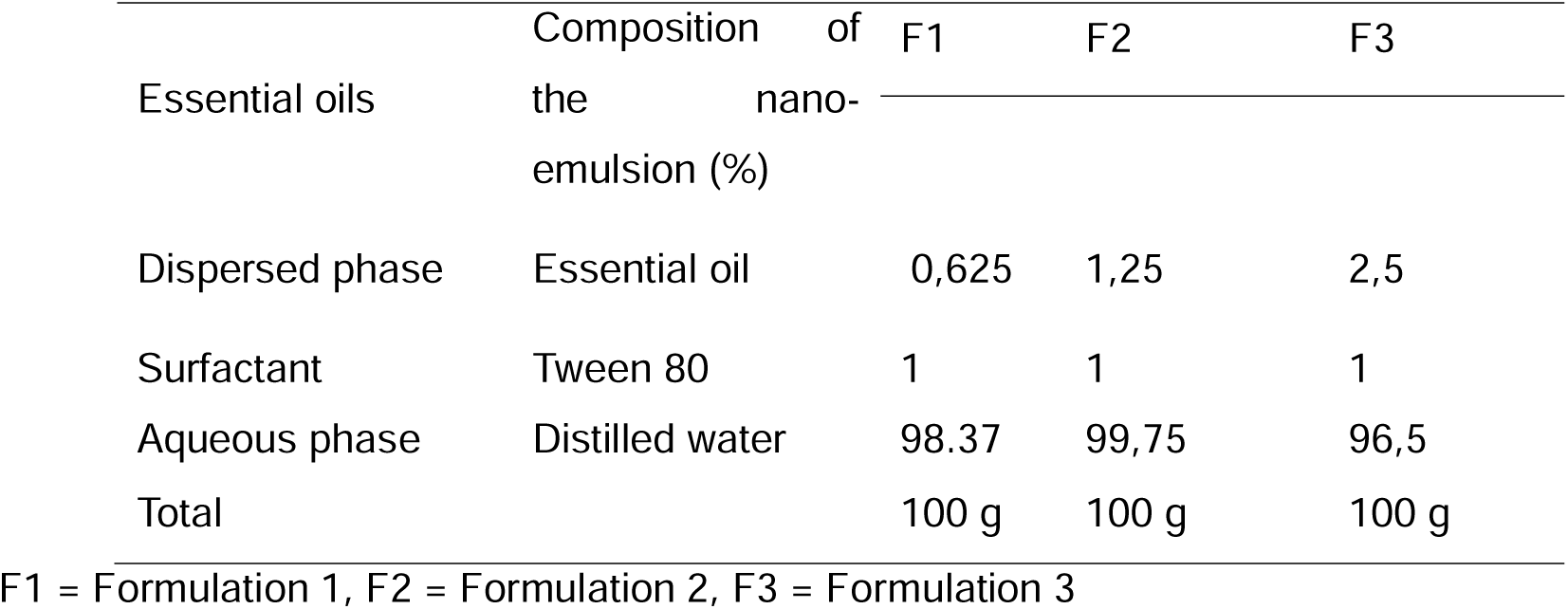
Formulation composition.

### Characterization of the nano-emulsion Stability

The stability of the nano-emulsion was evaluated according to Nirmala *et al*., with slight modifications. [16] Freshly prepared formulations were stored at 4°C, 25°C, and 37°C, and their macroscopic appearance was monitored every two days. Long-term physical stability was further assessed for one month after the addition of methylene blue to facilitate visualization of possible phase separation.

### Particle size analysis

Dynamic Light Scattering (DLS) measurements were performed using a Malvern Zetasizer Nano S equipped with a 633 nm He-Ne laser and low-volume disposable sizing cuvettes. Formulations F1, F2, and F3 were analyzed using water as dispersant (refractive index = 1.330; viscosity = 0.8872 cP) at 25.0°C. Each measurement consisted of three independent runs with an acquisition time of 120 s.

### Dermal safety of *O. gratissimum* nano emulsions

Acute dermal toxicity was evaluated according to OECD Test Guideline 402 [17]. Briefly, rats were weighed, individually identified, and randomly assigned to three experimental groups. Twenty-four hours before treatment, the dorsal area of each animal was carefully shaved, and food was withheld for 12 h while water remained available ad libitum. Distilled water, Tween 80 (2000 mg/kg body weight), or OGNe (2000 mg/kg body weight) was topically applied to the designated groups (n = 3 per group). Animals were observed continuously during the first 30 min and subsequently at 1, 2, 4, 8, 16, 24, and 48 h, followed by daily observations for 14 consecutive days. Clinical parameters included erythema, changes in mucous membranes, respiratory pattern, fur appearance, ocular alterations, tremors, convulsions, salivation, diarrhea, lethargy, sleep, and coma.

### Antimicrobial activity of *O. gratissimum* nano-emulsions

#### Determination of minimum inhibitory concentrations (MIC)

MIC values were determined following the Clinical and Laboratory Standards Institute (CLSI) guidelines [18] with slight modifications involving the use of resazurin as a viability indicator in a 96-well microplate assay [19]. Briefly, 80 μL of two-fold serial dilutions of the nanoemulsions prepared in nutrient broth were dispensed into each well, followed by the addition of 80 μL of bacterial suspension standardized to 1.5 × 10L CFU/mL. Sterility control wells contained culture medium only. Final nanoemulsion concentrations ranged from 1.625 to 100 μg/mL, whereas ceftriaxone concentrations ranged from 0.15625 to 100 μg/mL. Plates were incubated at 37°C for 24 h before the addition of 10 μL of resazurin solution and further incubation for 1 h. A color change from blue to pink indicated bacterial growth. The MIC was defined as the lowest concentration preventing the color change and maintaining the blue coloration. The most susceptible bacterial strain was subsequently selected for time-kill kinetic analysis.

#### Minimum bactericidal concentration (MBC)

The minimum bactericidal concentration (MBC) was determined from wells showing no visible bacterial growth during the MIC assay. Twenty-five microliters from each selected well were aseptically transferred into new wells containing 75 μL of Mueller-Hinton broth, resulting in a four-fold dilution of the nanoemulsion to eliminate residual inhibitory activity. Following incubation at 37°C for 24 h, the lowest concentration showing no bacterial regrowth was recorded as the MBC [20]. Positive controls consisted of culture medium, bacterial inoculum, and ceftriaxone, whereas negative controls contained only culture medium and bacterial inoculum. All experiments were performed in duplicate.

#### Determination of the MBC/MIC ratio

The MBC/MIC ratio was calculated to determine whether the nanoemulsion exerted bactericidal or bacteriostatic activity. According to accepted criteria, an MBC/MIC ratio ≤ 4 indicates bactericidal activity, whereas a ratio > 4 indicates a bacteriostatic effect [20].

#### Time-kill kinetics

Time-kill assays were performed only on the most susceptible bacterial strain, following the protocol previously described by Chahal et al. [21]. Nanoemulsion concentrations corresponding to 0.5×MIC, 1×MIC, and 2×MIC were prepared and inoculated with bacterial suspensions adjusted to 1.5 × 10L CFU/mL. Cultures were incubated at 37°C, and 0.5 mL aliquots were collected after 0, 2, 4, 6, 8, 12, and 24 h. Each aliquot was mixed with 50 μL of resazurin solution, incubated for 1 h, and fluorescence was measured using excitation and emission wavelengths of 560 and 590 nm, respectively. Measurements were performed in duplicate, and bacterial growth kinetics were expressed as log fluorescence versus incubation time [22].

### Wound healing activity

#### Excision wound model

Rats were weighed and anesthetized according to Rodent Anesthesia Protocol ANE-02 (December 6, 2000) by intraperitoneal administration of a ketamine/diazepam mixture (75 mg/kg and 5 mg/kg body weight, respectively) [23]. After shaving and disinfecting the dorsal skin with 70% ethanol, a full-thickness excisional wound measuring approximately 1 cm² was created using sterile surgical forceps and scissors. Animals were randomly allocated into five groups: Group 1, untreated negative control; Group 2, trolamine-treated positive control (400 mg/kg body weight); Group 3, OGNe containing 0.625% essential oil (400 mg/kg body weight); Group 4, OGNe containing 1.25% essential oil (400 mg/kg body weight); and Group 5, OGNe containing 2.5% essential oil (400 mg/kg body weight).

#### Evaluation of wound healing

Wound progression was monitored by measuring wound surface area every two days during the first 15 days and again on day 21 using a digital caliper. Measurements were continued until complete healing was achieved in the positive control group.

### Statistical analysis

Experimental data were recorded using Microsoft Excel 2013 and analyzed with GraphPad Prism version 10.6.0 (GraphPad Software, San Diego, CA, USA). Differences among groups were assessed by one-way or two-way analysis of variance (ANOVA), as appropriate, followed by Tukey’s multiple comparison test. Statistical significance was established at p < 0.05.

## Results

### Physicochemical characterization of *O. gratissimum* nanoemulsion

Hydrodistillation of fresh *O. gratissimum* leaves yielded 0.63% (w/w) essential oil. The resulting oil-in-water nanoemulsions exhibited a homogeneous milky-white appearance, a near-neutral pH (6.8), and persistent foam formation (Figure 2), indicating successful emulsification.

**Figure 2.**
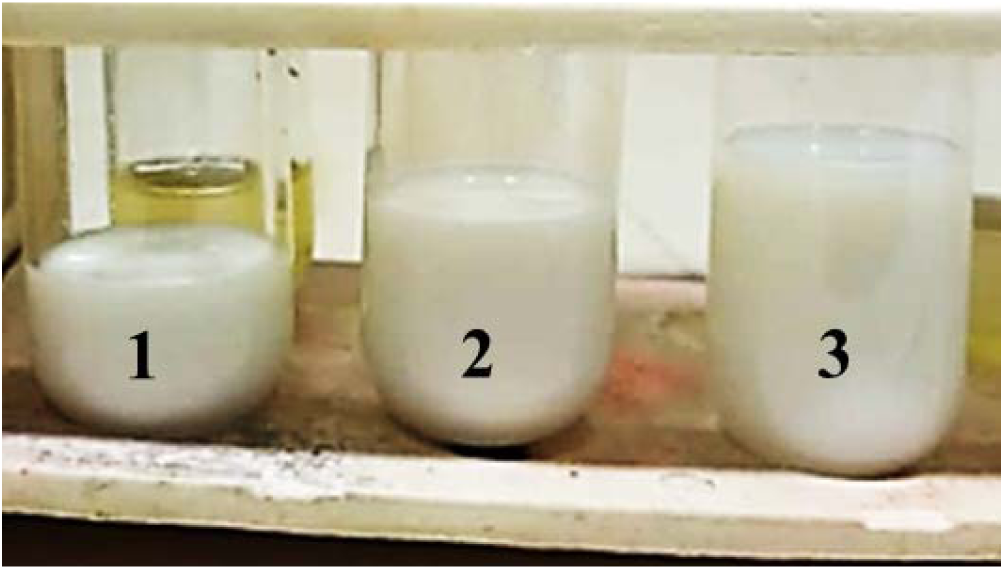
*O. gratissimum* essential oil and TWEEN 80 nano-droplets emulsion in water. 1 = OGNe 0,625 %, 2 = OGNe 1,25 %, 3 = OGNe 2,5 %,

### Stability

The physical stability of the formulated nanoemulsions was evaluated under different storage temperatures. After 14 days, the formulation stored at 37°C remained physically stable, retaining its homogeneous bluish appearance without evidence of phase separation or creaming. Similarly, the nanoemulsion stored at 25°C remained stable throughout the two-month observation period. Overall, the physicochemical characteristics of the formulations were maintained during storage (Table 3), demonstrating satisfactory stability under ambient and physiological temperature conditions. In contrast, the slight clarification observed after 14 days at 4°C may be attributed to temperature-induced changes in the aqueous continuous phase, which could affect light scattering without compromising the overall integrity of the nanoemulsion.

**Table 3.** Physicochemical characteristics and storage stability of *Ocimum gratissimum* nanoemulsions.

| Characteristics and stability tests | Results |
| --- | --- |
| Appearance of the essential oil | yellowish |
| Appearance after preparation | Milky |
| pH | 6.8 |
| Temperature stability after 14 days of storage |  |
| 4 °C | + |
| 25 °C | - |
| 37 °C | - |

- Absence of phase separation, + Low separation

### Dynamic light scattering

Dynamic light scattering analysis was performed to determine the particle size distribution, hydrodynamic diameter, and polydispersity of the three nanoemulsion formulations (F1, F2, and F3). The intensity- and volume-weighted distributions of formulations F1 and F3 exhibited multiple peaks, whereas formulation F2 displayed a more homogeneous profile, characterized by a predominant particle population in both the volume and number distributions. Importantly, all formulations fulfilled the instrument quality criteria, confirming the reliability of the measurements. The mean hydrodynamic diameters were 131.6 ± 91.8 nm, 139.7 ± 76.4 nm, and 69.5 ± 43.0 nm for F1, F2, and F3, respectively, while the corresponding polydispersity index (PDI) values were 0.487, 0.299, and 0.382. Similar autocorrelation (correlogram) profiles were obtained for all three formulations, indicating excellent measurement reproducibility (Figure 4A–C).

**Figure 4.**
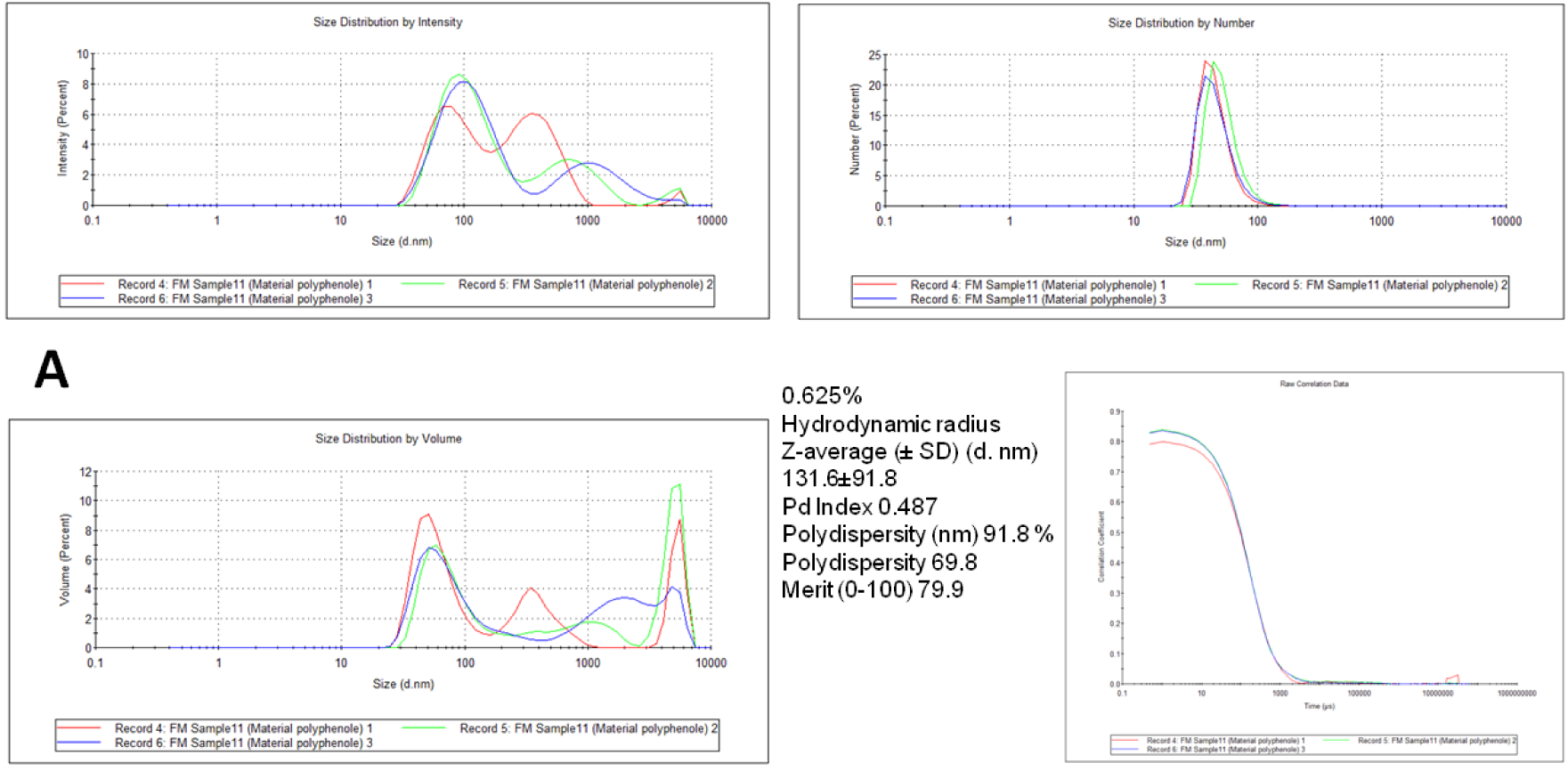
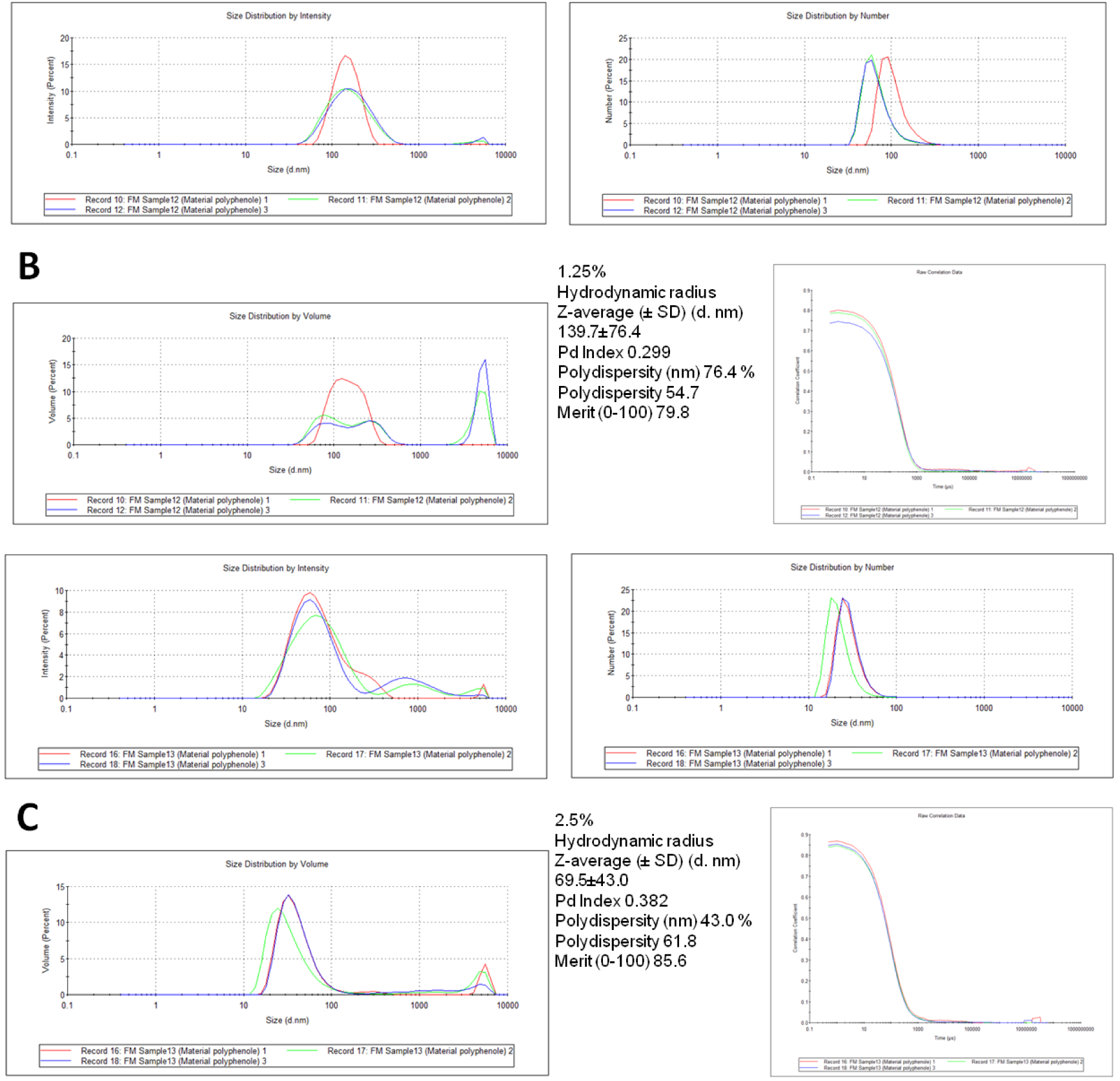
A) Dynamic light scattering characterization of *Ocimum gratissimum* nano-oil dispersions at three concentrations. (A) Size distribution profiles and correlation curves for the 0.625% formulation showing intensity-, number-, and volume-weighted particle size distributions from three independent measurements. (B) Size distribution profiles and correlation curves for the 1.25% nano-oil dispersion, illustrating changes in hydrodynamic diameter and polydispersity with increasing concentration. (C) Particle size distributions and autocorrelation data for the 2.5% formulation, highlighting the shift toward smaller nanoscale droplets and improved dispersion quality. All measurements were performed in triplicate to assess reproducibility.

The 0.625% nanoemulsion (Figure 4A) displayed a dominant nanosized particle population below approximately 100 nm together with a broader secondary population extending into the submicron range (approximately 400–2000 nm) in the intensity- and volume-weighted distributions. Because larger particles scatter light more efficiently, this secondary population likely represented a limited number of larger droplets contributing disproportionately to the scattering signal. The formulation exhibited a mean hydrodynamic diameter of 131.6 ± 91.8 nm and a PDI of 0.487, indicating moderate heterogeneity. The superimposition of the autocorrelation curves obtained from the three independent measurements confirmed the reproducibility of the analysis.

For the 1.25% formulation (Figure 4B), the particle size distribution became more compact and homogeneous, although the dominant population remained above 100 nm. The average hydrodynamic diameter was 139.7 ± 76.4 nm, whereas the PDI decreased to 0.299, reflecting improved dispersion uniformity compared with the 0.625% formulation. Although a small secondary population of larger droplets remained visible in the volume distribution, its contribution was markedly reduced. The close overlap of the correlograms further demonstrated the excellent stability of the dispersion during analysis.

Interestingly, the 2.5% formulation (Figure 4C) exhibited the smallest droplet size among all preparations. Both the intensity- and number-weighted distributions were centered predominantly below 100 nm, indicating efficient nanoemulsion formation. The mean hydrodynamic diameter decreased to 69.5 ± 43.0 nm, whereas the PDI increased slightly to 0.382, suggesting a moderate broadening of the particle size distribution despite the reduction in average droplet size. Only a negligible proportion of larger droplets was observed in the volume distribution. As with the other formulations, the autocorrelation curves showed excellent overlap between replicates, confirming high analytical reproducibility and colloidal stability.

Collectively, the DLS analysis demonstrated that increasing the essential oil concentration promoted the formation of smaller nanosized droplets, with the 2.5% formulation exhibiting the lowest average hydrodynamic diameter while maintaining satisfactory colloidal homogeneity and reproducibility.

### Acute dermal toxicity

The acute dermal toxicity study was conducted over a 14-day observation period following topical application of the nanoemulsions. No mortality was recorded in any treatment group, and none of the animals exhibited clinical signs of systemic or local toxicity. Specifically, no abnormalities in skin appearance, locomotor activity, pain sensitivity, respiratory pattern, ocular condition, salivation, convulsions, diarrhea, or behavioral responses were observed throughout the study (Table 4).

**Table 4.**
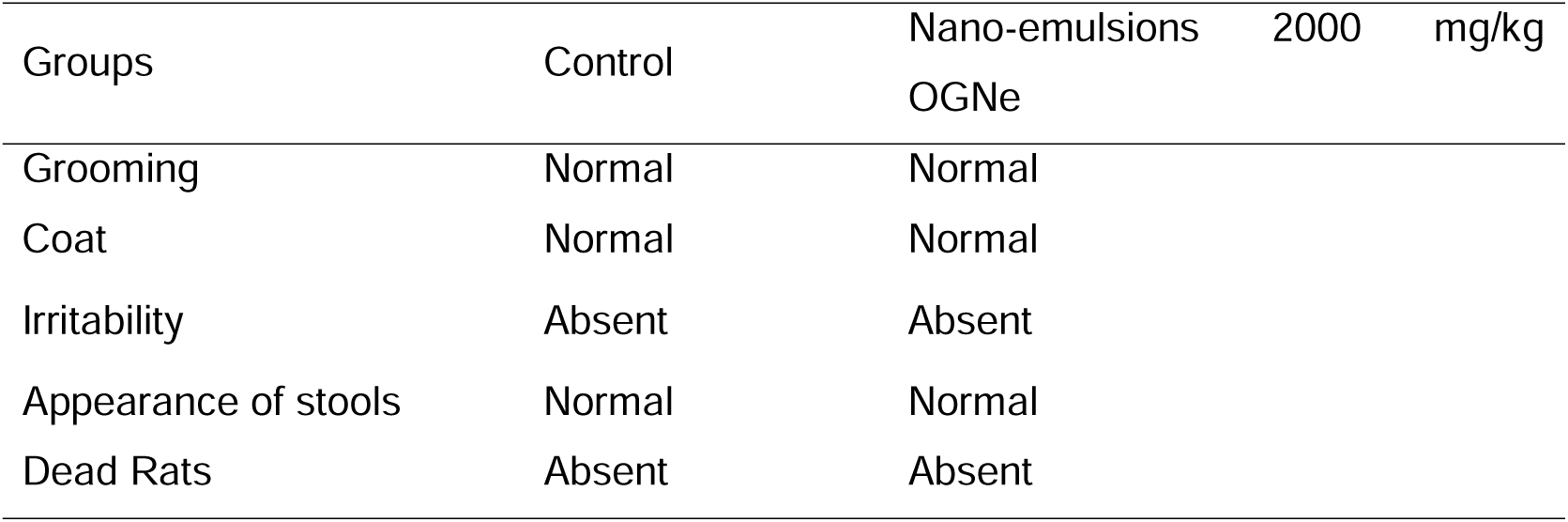
Clinical observations recorded during the acute dermal toxicity study in Wistar rats.

Body weight monitoring further confirmed the good dermal tolerance of the formulations (Figure 5). Animals treated with OGNe showed normal weight gain throughout the experimental period, comparable to that of the distilled water control group. Statistical analysis revealed a significant increase in body weight between days 1 and 11 in animals treated with the 1.25% formulation. Likewise, body weight at day 11 was significantly higher in the OGNe 1.25% group than in the control group. However, no significant differences were observed among the treated groups during the remainder of the study, indicating that topical administration of the nanoemulsions did not adversely affect normal growth or feeding behavior. These findings support the excellent dermal biocompatibility of the developed nanoemulsions.

**Figure 5:**
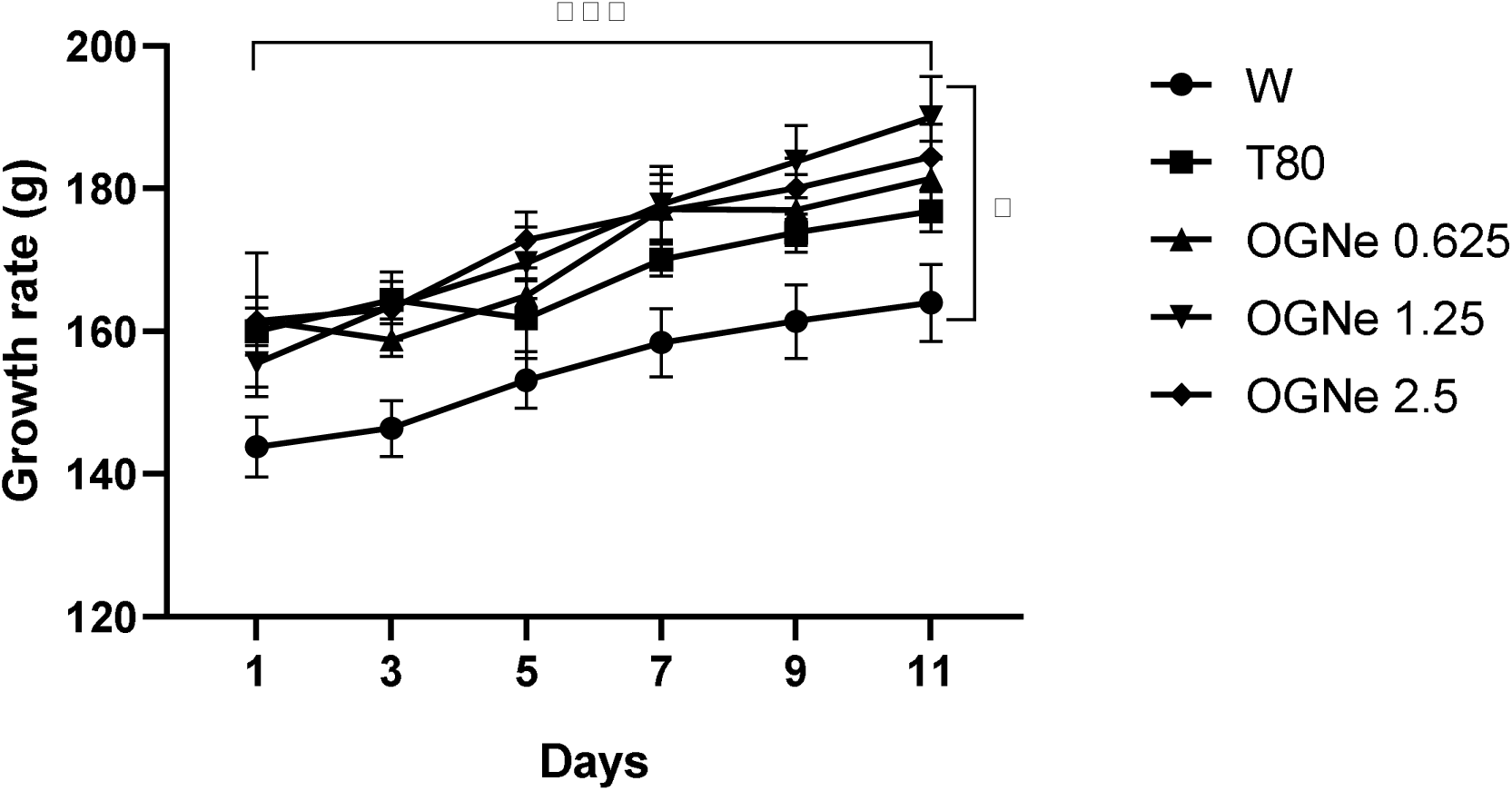
Evolution of the rat’s weight during the acute skin toxicity study. W = distilled water; T80 = Tween 80; OGNe = Nano-emulsion of essential oil of *O. gratissimum*

### Antibacterial activity of *Ocimum gratissimum* nanoemulsions

#### 1) Antibacterial activity

The antibacterial activity of the *Ocimum gratissimum* nanoemulsions (OGNe) was evaluated against selected bacterial pathogens using ceftriaxone as the positive control. The minimum inhibitory concentration (MIC) values obtained for each bacterial strain are summarized in Table 6.

**Table 6:** Minimum inhibitory concentrations of *Ocimum gratissimum* nanoemulsions against tested bacterial strains.

| Microorganisms | Ceftriaxone |  |  |
| --- | --- | --- | --- |
|  | MIC (µg/mL) | BMC(µg/mL) | BMC/MIC |
| *Acinetobacter baumannii* | 12.5 | 25 | 2 |
| *Escherichia coli* | 12,5 | 100 | 8 |
| *Pseudomonas aeruginosa* | 100 | 100 | 1 |
| *Proteus mirabilis* | 1,5625 | 1,5625 | 1 |

#### 2) Minimum inhibitory and bactericidal concentrations

The antibacterial efficacy of OGNe, expressed as MIC and MBC values, is presented in Table 7. The nanoemulsions inhibited the growth of all bacterial strains evaluated, with the exception of Escherichia coli, which remained resistant at all tested concentrations. Moderate inhibitory activity was observed against Proteus mirabilis and Pseudomonas aeruginosa, with MIC values of 6.25 μL/mL. The highest antibacterial susceptibility was recorded for Acinetobacter baumannii, for which the MIC was 1.25 μL/mL.

**Table 7.** Minimum inhibitory concentration (MIC), minimum bactericidal concentration (MBC), and MBC/MIC ratio of *Ocimum gratissimum* nanoemulsions.

| Isolates | MIC (en µL/mL ) | MBC (en µL/mL ) |
| --- | --- | --- |
| *Acinetobacter baumannii* | 1.125 | ND |
| *Escherichia coli* | ND | ND |
| *Proteus mirabilis* | 6.25 | ND |
| *Pseudomonas aeruginosa* | 6.25 | ND |

ND : Not determined

The enhanced activity observed against A. baumannii may be attributed to the presence of phenolic constituents, particularly eugenol, which is recognized for its membrane-disrupting and antimicrobial properties.

#### 3) Time-kill kinetics

The bactericidal kinetics of OGNe were investigated against the most susceptible bacterial strain, Acinetobacter baumannii, and are presented in Figure 6. At concentrations of 25 μL/mL and 12.5 μL/mL, bacterial counts remained nearly constant throughout the incubation period, indicating a predominantly bacteriostatic effect characterized by inhibition of bacterial multiplication rather than complete bacterial eradication. In contrast, treatment with 6.25 μL/mL produced only partial growth inhibition, as bacterial proliferation progressively increased over time, although remaining lower than that observed in the untreated control. These findings demonstrate a clear concentration-dependent antibacterial effect, with increasing OGNe concentrations producing stronger inhibition of bacterial growth.

**Figure 6:**
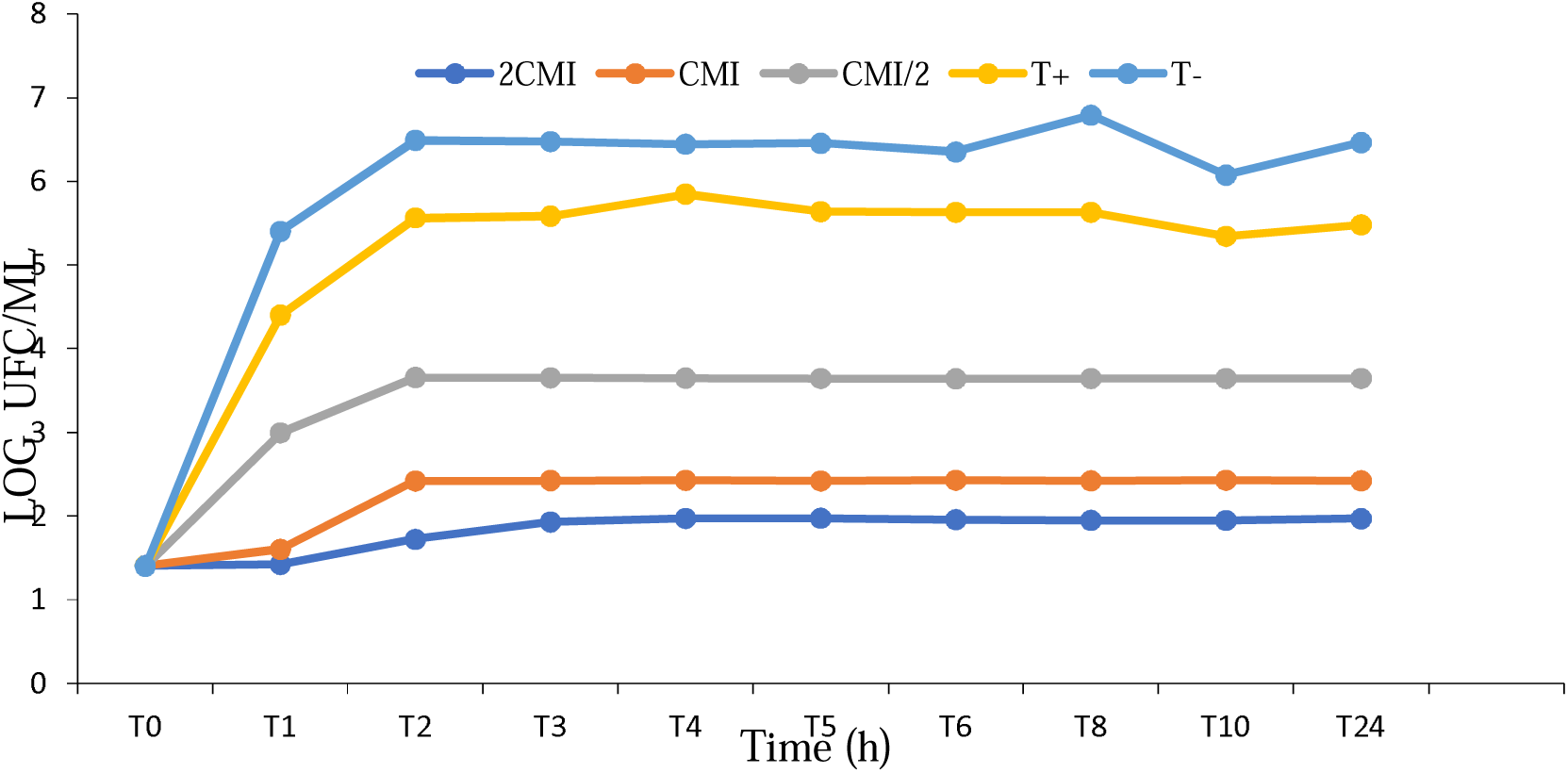
Mortality kinetics of *Acinetobacter baumanii* with OGNe

#### 4) Wound-healing activity

The wound-healing efficacy of *Ocimum gratissimum* nanoemulsions (OGNe) was evaluated using an excisional wound model in Wistar rats over a 15-day treatment period, with final healing assessed on day 21. Wound contraction was measured at predetermined intervals and compared with untreated animals receiving distilled water and with animals treated with the reference drug, trolamine. OGNe markedly accelerated wound closure during the inflammatory and proliferative phases of healing, demonstrating significantly greater contraction rates than the water-treated controls. By day 21, all groups exhibited nearly complete wound closure, and no significant differences were observed among the treated groups, indicating convergence toward full tissue repair. These findings suggest that OGNe enhances the early stages of wound healing without compromising the completion of the regenerative process (Figure 7).

**Figure 7:**
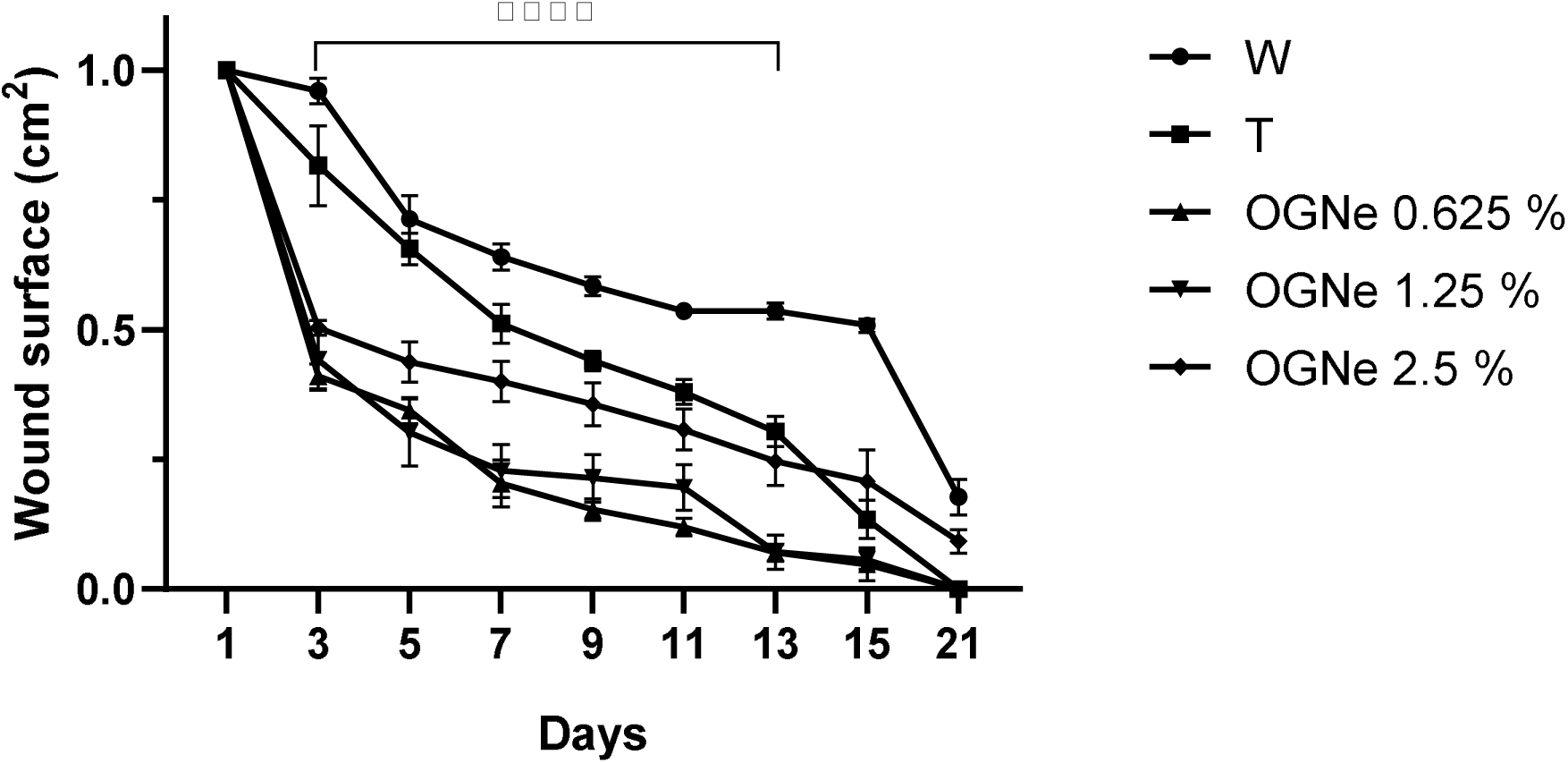
Time-course of wound surface closure in Wistar rats treated with water (W), trolamine (T, positive control), and *Ocimum gratissimum* nanoemulsions (OGNe; 0.625, 1.25, and 2.5 mg/kg). Wound contraction was monitored over 21 days and analyzed using Tukey’s multiple comparisons test. No significant differences were observed at day 1. From day 3 to day 13, OGNe-treated groups showed significantly greater wound closure compared with water- and trolamine treated controls (****p < 0.0001 at most time points). By day 21, wound closure rates converged among treated groups, indicating near-complete healing. The rate is significative only relative to water control treatments. Data are expressed as mean ± SEM. *p < 0.05, **p < 0.01, ***p < 0.001, ****p < 0.0001; ns: not significant.

## Discussion

The present study demonstrates that nanoemulsification is an effective strategy for improving the pharmaceutical properties of *Ocimum gratissimum* essential oil while preserving its biological activity. The formulation combined favorable physicochemical characteristics with good dermal tolerance, selective antibacterial activity, and accelerated wound repair, supporting its potential as a topical phytopharmaceutical.

The hydrodistillation yield (0.63%) was consistent with previous reports for *O. gratissimum* collected in the same geographical region, suggesting relatively consistent essential oil production under similar environmental conditions. Lower extraction yields reported from other Cameroonian localities emphasize the influence of geographical origin, climatic conditions, plant chemotype, harvesting season, and post-harvest handling on essential oil composition and recovery. [5, 8]

Physicochemical characterization confirmed the successful formation of stable oil-in-water nanoemulsions exhibiting a homogeneous milky appearance, near-physiological pH, and satisfactory storage stability. Such characteristics are desirable for topical formulations because they improve patient acceptability while minimizing the risk of skin irritation and preserving volatile phytoconstituents during storage.

Dynamic light scattering confirmed the formation of nanosized droplets with diameters ranging from approximately 26 to 224 nm. Although moderate polydispersity was observed, all formulations exhibited PDI values below 0.5, indicating acceptable dispersion homogeneity for mechanically emulsified nanoemulsions. Interestingly, the highest oil concentration generated the smallest average droplet size, suggesting more efficient stabilization of newly formed interfaces through improved surfactant coverage during emulsification rather than a simple concentration effect. Similar observations have been reported for plant essential oil nanoemulsions prepared by mechanical stirring or ultrasonic homogenization, where formulation composition largely determines droplet size and long-term stability. [24]

An important aspect of DLS interpretation concerns the discrepancy between intensity and number distributions. The secondary peaks observed in the intensity and volume profiles of the 0.625% and 1.25% dispersions, most likely correspond to minor aggregates rather than indicating poor formulation quality. The corresponding number distributions demonstrate that most droplets remained within the nanoscale domain, confirming efficient emulsification.

The antibacterial evaluation revealed selective activity against *Acinetobacter baumannii*, whereas *Escherichia coli* remained resistant. This finding is consistent with previous reports indicating that Gram-negative bacteria possessing highly impermeable outer membranes exhibit variable susceptibility to *O. gratissimum* essential oil. The observed bacteriostatic rather than bactericidal activity suggests that the nanoemulsion primarily interferes with bacterial proliferation instead of inducing immediate cell death. Such growth inhibition may nevertheless represent a considerable therapeutic advantage during wound healing by limiting bacterial expansion while allowing host immune mechanisms to eliminate residual microorganisms. *Acinetobacter baumannii* is increasingly recognized as an opportunistic pathogen associated with chronic wound infections and multidrug resistance; therefore, even moderate inhibition of its growth may have clinical relevance. [6, 25] The bacteriostatic activity demonstrated by the time-kill assay indicates that OGNe inhibits bacterial multiplication rather than causing rapid bacterial death. Such growth suppression may nevertheless be therapeutically advantageous, allowing host immune defenses sufficient time to eliminate residual microorganisms while reducing bacterial colonization at the wound site. [25]

Topical administration produced no evidence of dermal irritation, behavioral abnormalities or systemic toxicity. This safety profile agrees with the long history of culinary and medicinal use of *O. gratissimum*, whose essential oil has previously demonstrated low toxicity in experimental models. [6] Encapsulation within nanoemulsions may further improve local tolerance by controlling the release of volatile constituents and reducing direct exposure of the skin to highly concentrated essential oil components. The wound-healing evaluation demonstrated that OGNe significantly accelerated wound contraction during the inflammatory and proliferative phases compared with untreated animals. Rapid wound contraction during these early stages is particularly important because it shortens tissue exposure, reduces the likelihood of microbial colonization, and limits excessive inflammation. Although all treated wounds reached nearly complete closure by day 21, the early acceleration of healing produced by OGNe represents a clinically meaningful advantage.

The polydispersity index values (0.299-0.487) observed across the formulations suggest moderately polydisperse systems, which is typical for nanoemulsions produced through conventional emulsification techniques. The intermediate concentration (1.25%) showed the lowest PDI, indicating the most uniform droplet distribution among the three formulations. However, despite the slightly higher PDI at 2.5%, the overall droplet size remained significantly smaller, which may provide advantages for certain applications. Droplet sizes below 100 nm, such as those observed for the 2.5% formulation, are particularly advantageous for enhancing interfacial surface area and improving the diffusion of encapsulated bioactive compounds. In essential oil nanoemulsions, smaller droplet sizes provide a larger interfacial surface area that enhances colloidal stability, promotes interactions with biological membranes, and improves the antimicrobial efficacy of encapsulated phytochemicals. [26] The nanoscale droplets observed here therefore suggest promising potential for biological or pharmacological applications involving *O. gratissimum* oil.

Furthermore, the strong overlap of the autocorrelation curves across replicate measurements indicates excellent reproducibility and colloidal stability during the analysis, confirming that the prepared dispersions maintain consistent scattering behavior.

Taken together, these results indicate that increasing the concentration of *O. gratissimum* nano-oil influences both the particle size distribution and dispersion homogeneity, with the 2.5% formulation producing the smallest nanoscale droplets, while the 1.25% formulation displays the most uniform distribution. These findings provide valuable insight for optimizing nano-oil formulations intended for biological or therapeutic applications.

Nanoemulsification likely potentiates the biological properties by increasing the contact surface between the oil droplets and injured tissue, facilitating penetration into superficial skin layers and improving the local bioavailability of active constituents.

The wound healing effect of the nano-emulsion was dose dependent. The *O. gratissimum* essential oil nano-emulsion comparison with the positive control, trolamine justifies its use in the nano-formulation of future natural healing.

Dose-response comparisons revealed modest differences among OGNe concentrations, although the 0.625 and 1.25 mg/kg formulations often produced statistically signifiant closure rates than the 2.5 mg/kg dose, reaching statistical significance at several time points (days 5–15). This observation suggests that maximal biological activity is not necessarily associated with the highest essential oil concentration but rather with an optimal balance between droplet size, formulation homogeneity, release kinetics, and local tissue bioavailability. Similar non-linear dose-response relationships have previously been described for phytochemical-loaded nanoemulsions intended for topical delivery. Importantly, by the late healing phase (day 21), differences between T and OGNe treatments were no longer significant, suggesting convergence toward complete tissue repair. Nonetheless, early-phase acceleration of wound contraction remains clinically relevant, as rapid closure reduces infection risk and inflammatory burden.

The data indicate that *O. gratissimum* nanoemulsions significantly enhance early and intermediate wound healing compared with untreated controls and show comparable or superior performance to the standard topical reference. [27] The enhanced healing probably results from the synergistic combination of several mechanisms. Nanoemulsification increases the contact surface between the encapsulated essential oil and injured tissue, promotes penetration through damaged skin, prolongs local retention of bioactive compounds, and enhances antioxidant and antimicrobial activities. Together, these effects create a microenvironment favorable for fibroblast proliferation, collagen deposition, angiogenesis, and tissue remodeling, thereby accelerating wound repair. These findings align with recent literature reporting that nanoemulsion-based essential oils improve dermal penetration, antioxidant activity, and local antimicrobial effects, collectively promoting faster tissue regeneration and collagen remodeling. [28]

## Conclusion

This study demonstrates that nanoemulsification is an efficient strategy for improving the pharmaceutical performance of *Ocimum gratissimum* essential oil while preserving its intrinsic biological activity. The developed oil-in-water nanoemulsions exhibited favorable physicochemical properties, including nanometric droplet size, acceptable homogeneity, near-physiological pH, and satisfactory storage stability, confirming the successful incorporation of the essential oil into a stable topical delivery system.

Topical administration produced no evidence of acute dermal toxicity, skin irritation, or behavioral abnormalities, supporting the excellent biocompatibility of the formulation. Antibacterial evaluation revealed selective inhibitory activity, particularly against *Acinetobacter baumannii*, whereas time-kill assays demonstrated a concentration-dependent bacteriostatic effect. Although *Escherichia coli* remained resistant, the antimicrobial profile observed is consistent with the known mechanisms of phenolic phytoconstituents, whose efficacy may be enhanced by nanoemulsion-mediated interactions with bacterial membranes. The most remarkable finding of this work was the significant acceleration of wound contraction throughout the inflammatory and proliferative phases of tissue repair. Compared with untreated animals, OGNe promoted faster wound closure and achieved healing outcomes comparable to, or exceeding, those obtained with the reference treatment, trolamine. The improved therapeutic performance is likely attributable to enhanced dispersion of the essential oil, increased dermal penetration, prolonged local retention of bioactive constituents, and the combined antimicrobial and antioxidant properties of the encapsulated phytochemicals.

Collectively, the physicochemical characterization, dermal safety assessment, antibacterial evaluation, time-kill kinetics, and wound-healing experiments provide compelling evidence that *Ocimum gratissimum* nanoemulsions constitute a promising phytopharmaceutical platform for topical wound management. Future studies should investigate release kinetics, collagen synthesis, angiogenesis, inflammatory mediators, and histopathological changes in both sterile and infected wound models to further elucidate the mechanisms underlying tissue regeneration and facilitate future clinical translation.

## Author contribution

Fomesseng Negoue A Investigation, animal models, writing - original draft; writing - review & editing. Eya’ane Meva F Conceptualization, supervision, DLS investigation, writing - original draft; writing - review & editing Hzounda Fokou J B, Methodology, software, validation, supervision, microbiology, writing - original draft; writing - review & editing Voundi Olugu S, Methodology, stability, validation, writing - review & editing. Boudjeka V Methodology, stability, validation, writing - review & editing, Ngo Nyobe J C Methodology, validation, writing - review & editing, Belle Ebanda Kedi P Methodology, animal models validation, writing - review & editing, Houatchaing Kouemegne A M Methodology, validation, DLS interpretation, writing - review & editing, Etame Loe G Conceptualization, supervision, writing - review & editing

## Declarations

## Data availability

The authors declare that the data supporting the findings of this study including raw data files are available from the corresponding author upon reasonable request.

## Statement on animal rights

On behalf of all authors, the corresponding author affirms that animal rights were upheld in the study. Ethics approval Ethical clearance Nr. 3580CEI-UDo/05/2023/T was obtained from the Institutional Research Ethics Committee for Human Health of the University of Douala (CEI-UDo).

